# Evidence of stock connectivity, hybridization and misidentification in white anglerfish support the need of a genetics-informed fisheries management framework

**DOI:** 10.1101/2021.02.10.430581

**Authors:** Imanol Aguirre-Sarabia, Natalia Díaz-Arce, Iker Pereda-Agirre, Iñaki Mendibil, Agurtzane Urtizberea, Hans D. Gerritsen, Finlay Burns, Ian Holmes, Jorge Landa, Ilaria Coscia, Iñaki Quinconces, Marina Santurtún, Antonella Zanzi, Jann T. Martinsohn, Naiara Rodríguez-Ezpeleta

## Abstract

Understanding population connectivity within a species as well as potential interactions with its close relatives is crucial to define management units and to derive efficient management actions. However, although genetics can reveal mismatches between biological and management units and other relevant but hidden information such as species misidentification or hybridization, the uptake of genetic methods by the fisheries management process is far from having been consolidated. Here, we have assessed the power of genetics to better understand the population connectivity of white angelfish (*Lophius piscatorius*) and its interaction with its sister species, the black anglerfish (*L. budegassa*). Our analyses, based on thousands of genome-wide single nucleotide polymorphisms, show three findings that are crucial for white anglerfish management. We found i) that white anglerfish is likely composed of a single panmictic population throughout the Northeast Atlantic, challenging the three-stock based management, ii) that a fraction of specimens classified as white anglerfish using morphological characteristics are genetically identified as black anglerfish (*L. budegassa*) and iii) that the two *Lophius* species naturally hybridize leading to a population of hybrids of up to 20% in certain areas. Our results set the basics for a genetics-informed white anglerfish assessment framework that accounts for stock connectivity, revises and establishes new diagnostic characters for *Lophius* species identification and evaluates the effect of hybrids in the current and future assessments of the white anglerfish. Furthermore, our study contributes to provide additional evidence of the potentially negative consequences of ignoring genetic data for assessing fisheries resources.

## Introduction

Sustainable exploitation of fisheries resources relies on effective fisheries management actions, which in turn rely on accurate fisheries assessment, that is, the process that includes the synthesis of information on life history, fishery monitoring, and resource surveys for estimating stock size and harvest rate relative to sustainable reference points (Methot & Wetzel, 2013). The process of fisheries assessment is usually applied independently to pre-established management units (so-called stocks), of which parameters such as growth, recruitment, and natural and fishing mortality are assumed to be intrinsic and not dependent on emigration or immigration rates (Cadrin, 2020). Genetic data has demonstrated ability to delineate populations within a species, that is, to identify groups of sexually interbreeding individuals which possess a common gene pool, but has also revealed hidden phenomena such as species misidentification (Garcia-Vazquez, Machado-Schiaffino, Campo, & Juanes, 2012) or hybridization (Pampoulie et al., 2020).

Fisheries stocks are often defined upon political and administrative considerations (Stephenson, 2002); yet, failure to align stocks with natural populations results in unfeasibility to establish an accurate relationship between productivity and harvest rates and can result in local reduction of populations and, in extreme cases, to local population collapse (Bonfil, 2005). Misidentification can be a common phenomenon when morphologically similar species are caught simultaneously, and results in misled exploitation rate estimations when those are based on reported catches (Marko et al., 2004). Hybridization, the successful reproduction between different species (Arnold, 1997; Stronen & Paquet, 2013), has been reported in teleost fishes (Schwenke, 2013; Yaakub, Bellwood, van Herwerden, & Walsh, 2006), but incidence and associated consequences in commercial fisheries has been scarcely explored (Epifanio & Nielsen, 2000); yet, hybridization could play a key role in diversity loss and even parental species extinction, which has important consequences for management and conservation (Allendorf, Leary, Spruell, & Wenburg, 2001). An example of species whose management could largely benefit from genetic-derived information is the white anglerfish (*Lophius piscatorius*, Linnaeus, 1758), a bottom-living fish that moved from being a bycatch species in the last century (Hislop et al., 2001) to become one of the most valuable demersal species in southern and western Europe (ICES, 2010).

The white anglerfish inhabits the Northeast Atlantic and Mediterranean Sea, where it is assessed by the International Council for the Exploration of the Sea (ICES) and the General Fisheries Commission for the Mediterranean (GFCM), respectively. Within the Northeast Atlantic, the white anglerfish is managed as three stocks (Figure 1): the Northern Shelf stock (Skagerrak, Kattegat, North Sea, West of Scotland and Rockall), the Northern stock (Celtic Seas and Northern Bay of Biscay) and the Southern Stock (Atlantic and Iberian waters) (ICES, 2019a, 2019b); yet, the few studies assessing the population structure, based on otolith shape analysis (Cañás, Stransky, Schlickeisen, Sampedro, & Fariña, 2012), tagging surveys (J. Landa, Quincoces, Duarte, Fariña, & Dupouy, 2008; Laurenson, Johnson, & Priede, 2005) and molecular markers, including allozymes (Crozier, 1987), mitochondrial DNA (Charrier et al., 2006) and microsatellites (Blanco, Borrell, Cagigas, Vázquez, & Prado, 2008), did not find differences between stocks. However, this needs to be confirmed with the analysis of a large number of genomic markers, which has been effective in resolving the population structure of marine fish when other markers failed (Leone, Álvarez, García, Saborido-Rey, & Rodriguez-Ezpeleta, 2019; Rodríguez-Ezpeleta et al., 2016; Rodríguez-Ezpeleta et al., 2019). The white anglerfish coexists with its sympatric sister species, the black bellied anglerfish (*Lophius budegassa*), which has a more southern distribution (Relini, 1999; Ungaro et al., 2002). The outer morphology of both species is similar which is why they are often confused, although the color of the epithelium that covers the abdominal cavity, called peritoneum (white in *L. piscatorius* and black in *L. budegassa*) is considered a unequivocal diagnostic character for individuals larger than 15 cm (Caruso, 1986). Yet, genetic analyses based on polymerase chain reaction amplification of restriction fragment length polymorphisms (PCR-RFLP) used for species identification have revealed mislabeling among the two species (Armani et al., 2012).

**Figure 1.**
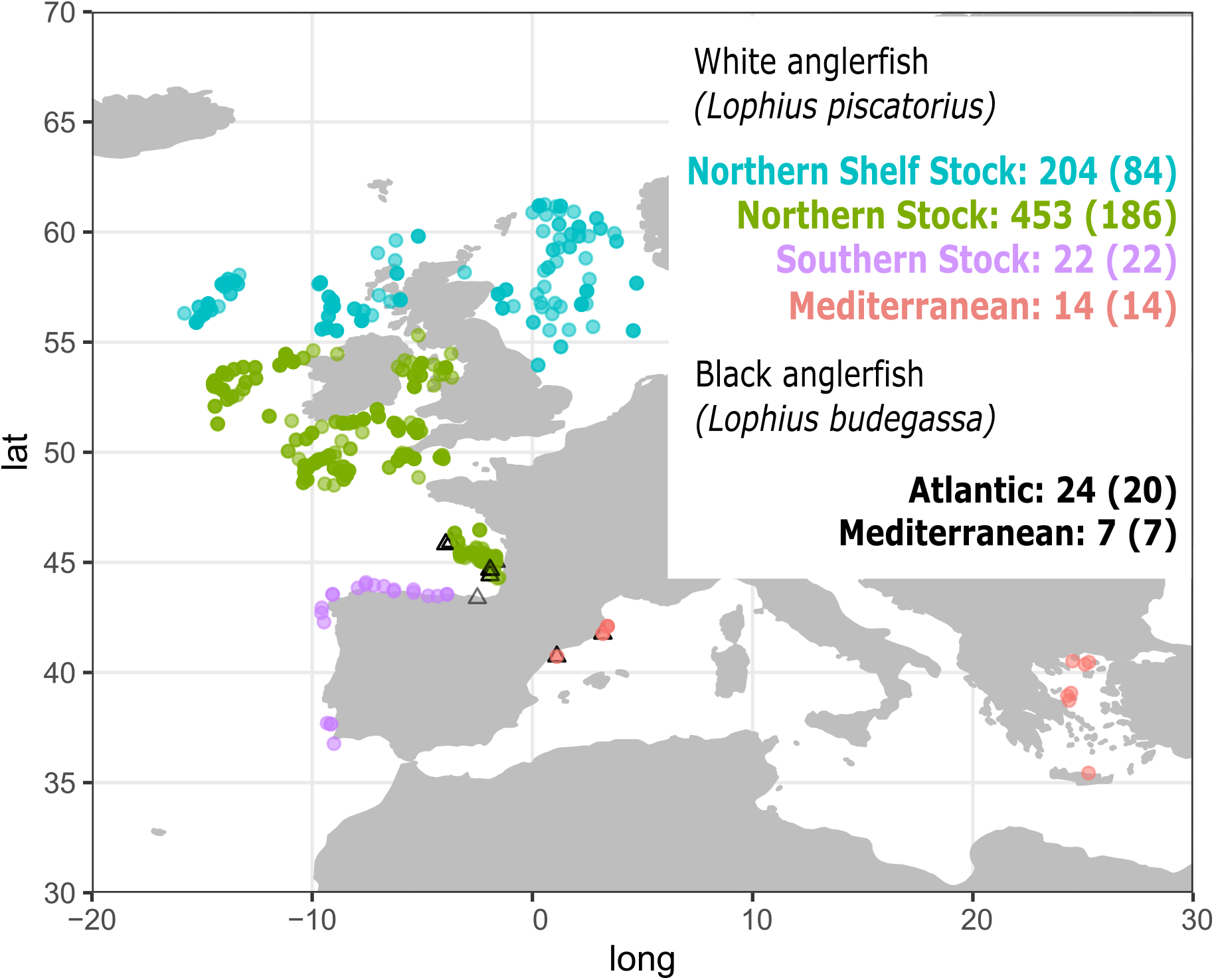
Samples included in this study, and in parentheses those genotyped with RAD-seq, where black triangles indicate Lophius budegassa samples and coloured circles, Mediterranean (red), Atlantic southern stock (purple), Atlantic northern stock (green) or Atlantic northern shelf stock (blue) Lophius piscatorius samples.

There is thus a need for resolving the population structure of white anglerfish within the Northeast Atlantic, including its relationship with Mediterranean populations, and for understanding its interaction with its sister species, the black anglerfish. For that aim, we have used thousands of genome-wide distributed SNP markers discovered and genotyped through restriction site associated DNA sequencing (RAD-seq) and found i) that white anglerfish from the three stocks within the Northeast Atlantic are genetically connected, ii) first evidence of hybridization between white and black anglerfish and iii) records of black anglerfish misidentified as white anglerfish due to a lack of black peritoneum. These findings have important implications for white anglerfish management and conservation, while revealing the significant advantage of including genomics in fisheries assessment in general.

## Material and methods

### Tissue sampling, DNA extraction, and species identification

*Lophius piscatorius* (n=693) and *Lophius budegassa* (n=31) samples were collected from Northeast Atlantic Ocean and Mediterranean Sea locations using scientific surveys and commercial fisheries (Table S1; Figure 1). Sampling of *L. piscatorius* was carried out so as to cover a large part of the geographic range of this species within the Atlantic, whereas samples of *L. budegassa* were collected opportunistically as they were only used for comparison purposes. Each individual was assigned to either species by the color of the peritoneum, white for *L. piscatorius* and black for *L. budegassa* (Caruso, 1986). Maturity was assigned following the 5 stages key (ICES, 2007). From each individual, a ∼1 cm^3^ muscle tissue sample was excised and immediately stored in 96% molecular grade ethanol at -20 °C until DNA extraction.

Genomic DNA was extracted from about 20 mg of muscle tissue using the Wizard® Genomic DNA Purification kit (Promega, WI, USA) following the manufacturer’s instructions. Extracted DNA was suspended in Milli-Q water and concentration was determined with the Quant-iT dsDNA HS assay kit using a Qubit® 2.0 Fluorometer (Life Technologies). DNA integrity was assessed by electrophoresis, migrating about 100 ng of Sybr™ Safe-stained DNA on an agarose 1.0% gel. A polymerase chain reaction restriction fragment length polymorphism (PCR-RFLP) method based on mitochondrial DNA (Armani et al., 2012) was used for authentication of all specimens collected. In order to further validate the PCR-RFLP results, for 122 of the samples, a fragment of the mitochondrial cytochrome b (cytb) gene was amplified with the primers GluFish-F and THR-Fish-R (Sevilla et al., 2007) using the following amplification conditions: denaturation at 95 °C for 3 min; 35 cycles at 95 °C for 30 s, 57 °C for 30 s, 72 °C for 60 s; and final extension at 72 °C for 10 min. The PCR products were purified and sequenced using Sanger.

### RAD-seq library preparation and sequencing

Restriction-site-associated DNA libraries were prepared for 306 *L. piscatorius* and 27 *L. budegassa* individuals (Table S1) following the methods of Etter, Bassham, Hohenlohe, Johnson, and Cresko (2012). Five individuals were run by duplicate starting from the tissue sample to check for replicability. Between 300 and 600 ng of starting DNA (depending on integrity) was digested with the *SbfI* restriction enzyme and ligated to modified Illumina P1 adapters containing 5bp sample-specific barcodes. Pools of 32 individuals were sheared using the Covaris® M220 Focused-ultrasonicator™ Instrument (Life Technologies) and size selected to 300-500 bp with magnetic beads. Following the Illumina P2 adaptor ligation, each library was amplified using 12 PCR cycles and batches of three pools were paired-end sequenced (100 bp) on an Illumina HiSeq4000.

### SNP genotype table generation

Generated RAD-tags were analyzed using Stacks version 2.4 (Catchen, Hohenlohe, Bassham, Amores, & Cresko, 2013). Quality filtering and demultiplexing were per-formed using the *Stacks* module *process_radtags* removing reads with ambiguous ba-ses and with an average quality score lower than 20 in at least one stretch of 15 nucle-otides and using a maximum of 1 mismatch when rescuing single-end barcodes. Only reads whose forward and reverse pair passed quality filtering were kept and the mod-ule *clone_filter* was applied to remove PCR duplicates. *De novo* and reference-based assemblies were performed for consistency since each approach has disadvantages: the genome used for the reference-based approach is highly fragmented (Dubin, Jørgensen, Moum, Johansen, & Jakt, 2019) and represents only one of the species in-cluded in this study, and the de novo approach could be affected by assembly parameters choice. For the *de novo* assembly, the module *ustacks* was used to assemble filtered reads into putative orthologous loci per individual, with a minimum coverage depth required to create a stack (parameter -m) of 3, and a maximum nucleotide mismatches allowed between stacks (parameter -M) of 4. RAD-loci were then assembled using the module *cstacks* with a maximum of 6 allowed mismatches between sample loci when generating the catalogue (parameter -n). Matches to the catalogue for each sample were searched using *sstacks* and transposed using *tsv2bam*. For the reference mapped assembly, the filtered reads were mapped against the available draft white anglerfish genome (Dubin et al., 2019) using the BWA-MEM algorithm (Li, 2013), and the resulting SAM files converted to BAM format, sorted and indexed using SAMTOOLS (Li et al., 2009). Mapped reads were filtered to include only primary alignments and correctly mate mapped reads. The following steps were applied to both the *de novo* and reference mapped catalogs including only samples with a minimum of 30,000 RAD-loci. Paired-end reads were assembled into contigs and SNPs derived were identified and genotyped using the module *gstacks*. In order to avoid ascertainment bias (Rodríguez-Ezpeleta et al., 2016), SNP selection was performed separately on each subset of individuals to be analyzed (Table S2). For that aim, the module *populations* was used to select the tags present in RAD-loci found in at least 90% of the individuals, and *PLINK* version 1.07 (Purcell et al., 2007) was used to select samples with a minimum of 0.85 genotyping rate, and SNPs with a minimum of 0.95 genotyping rate and a minimum allele frequency (MAF) bigger than 0.05. Tags on which the selected SNPs were located were mapped against the complete mitochondrial genome of *Lophius piscatorius* (NC_036988.1) with the BWA-MEM algorithm (Li, 2013) and it was con-firmed that the final genotype dataset did not contain mitochondrial SNPs.

### Genetic diversity, population structure and hybridization analyses

The following analyses were performed using only the first SNP per tag. Expected (He) and observed (Ho) heterozygosity and average pairwise F_ST_ values were computed with PLINK (Purcell, 2009) and GENEPOP (Rousset, 2014) respectively. Principal component analysis (PCA) was performed without any *a priori* assignment of individuals to populations using the package *adegenet* (Jombart & Ahmed, 2011) in R version 3.5.0 (Team, 2018). The genetic ancestry of each individual was estimated using the model-based clustering method implemented in ADMIXTURE (Alexander, Shringarpure, Novembre, & Lange, 2015) assuming from 2 to 5 ancestral populations (K) and setting 1000 bootstrap runs. The value of K with the lowest associated error value was identified using ADMIXTURE’s cross-validation procedure assuming from 1 to 5 ancestral populations. Using *NewHybrids* (E. Anderson & Thompson, 2002), posterior probabilities of each individual’s membership as a pure parent, first (F1) or second (F2) generation hybrids or backcrosses were calculated applying default parameters to the 200 SNPs with less than 1% of missing data that presented the highest Fst between genetically assigned *L. piscatorious* and *L. budegassa* samples.

## Results

### Genetic species identification based on mitochondrial markers

In all samples for which it was conducted, cytb gene sequencing confirmed PCR-RFLP results (Table S1). Species authentication was positive for all 31 specimens identified as *Lophius budegassa*. However, from the 693 specimens identified as *L. piscatorius* based on the colour of their peritoneum, 76 were genetically identified as *L. budegassa*. This incongruency between the colour of the peritoneum and the mitochondrial based genetic identification (Figure 2) suggests that species identification based on the colour of the peritoneum results in erroneous taxonomic assignment or that mitochondrial markers are ambiguous for species identification, which needs to be evaluated using nuclear markers.

**Figure 2.**
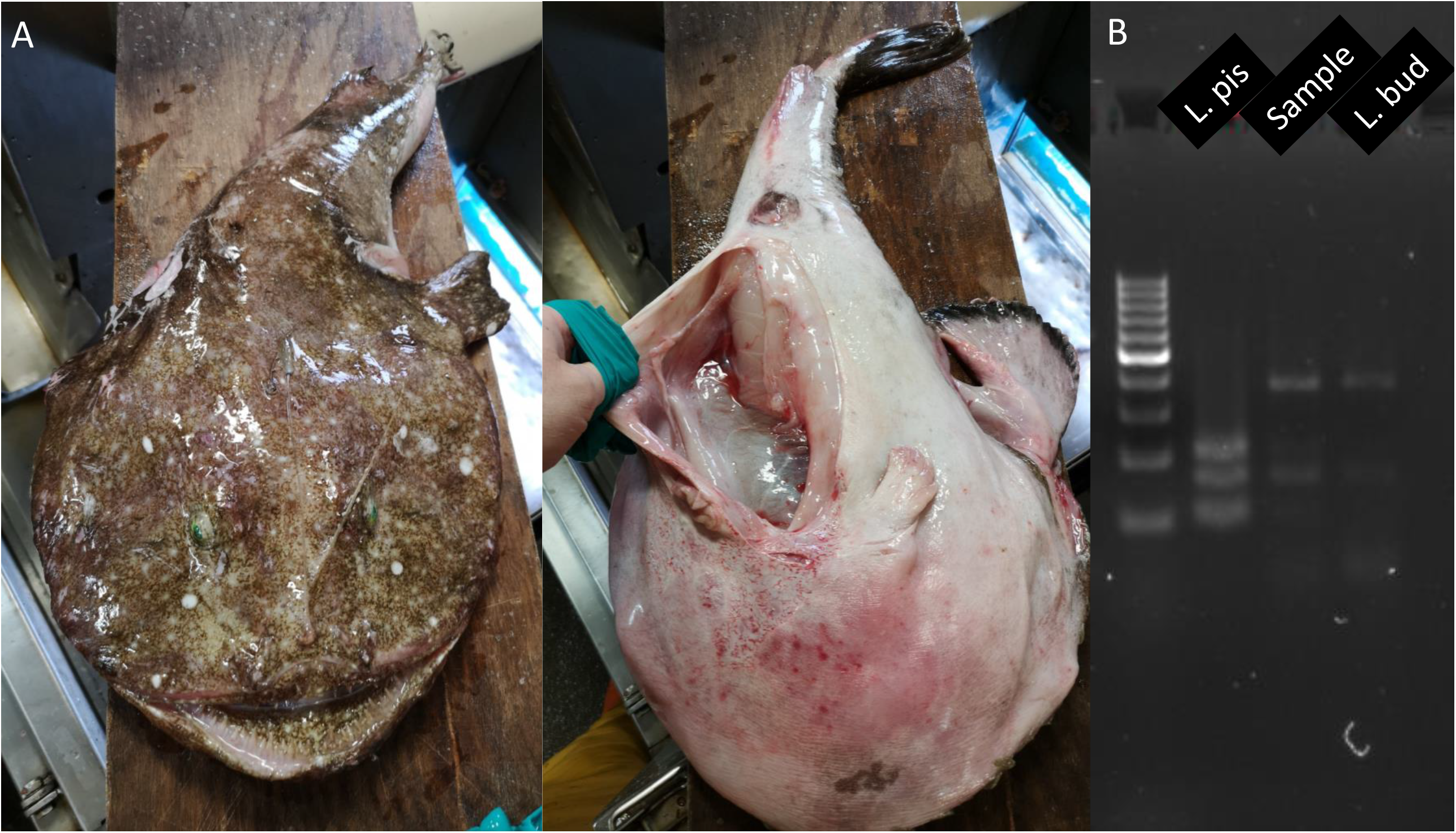
A. Specimen of anglerfish captured in the northern coast of Scotland (65 cm and 4.4 kg) with white peritoneum, emerald green eyes, and stippled skin, genetically identified as black anglerfish. B. Agarose gel electrophoresis of the resulting PCR-RFLP band for white (L. pis) and black anglerfish (L. bud) showing that this specimen (Sample) is genetically identified as black anglerfish.

### Population structure based on nuclear markers

The genotype table including individuals from both species resulted in 323 and 326 individuals (of which 27 were *L. budegassa*) and 16712 or 23126 SNPs after filtering when using a *de novo* or reference mapped assembly respectively (Table S2). Replicates resulted in 0.9947% identical genotypes. The PCA and ADMIXTURE analyses based on these datasets grouped most samples in three clearly distinguishable groups, and results were virtually identical among the *de novo* and reference assembly datasets (Figure 3; Figure S1) and subsequent analyses were based on the former. In the PCA (Figure 3A), samples are disposed along the first principal component (accounting for 80.82% of the total variance) into three main groups. The first group includes individuals provided, and genetically confirmed, as *L. piscatorius*. The second group (in the middle) includes 25 individuals initially identified as *L. piscatorius*, although 22 of them are assigned to *L. budegassa* according to mitochondrial DNA. The third group includes individuals initially identified as *L. piscatorius* and *L. budegassa*, but all of them assigned to *L. budegassa* according to mitochondrial DNA. Additionally, there are five individuals (all with *L. budegassa* mitochondrial DNA) located in between the main three clusters. This sample grouping is coherent with the different types of genetic admixture patterns observed according to the presence of two ancestral populations, best K=2 (Figure 3B): samples included in the first and third groups in the PCA are not admixed and belong to different ancestral populations, and samples included in the middle group in the PCA are admixed with equal contribution from both ancestral populations. Samples located between groups in the PCA show consistent patterns in admixture analyses with about 75% or 25% of contribution from the first and third groups, respectively.

**Figure 3.**
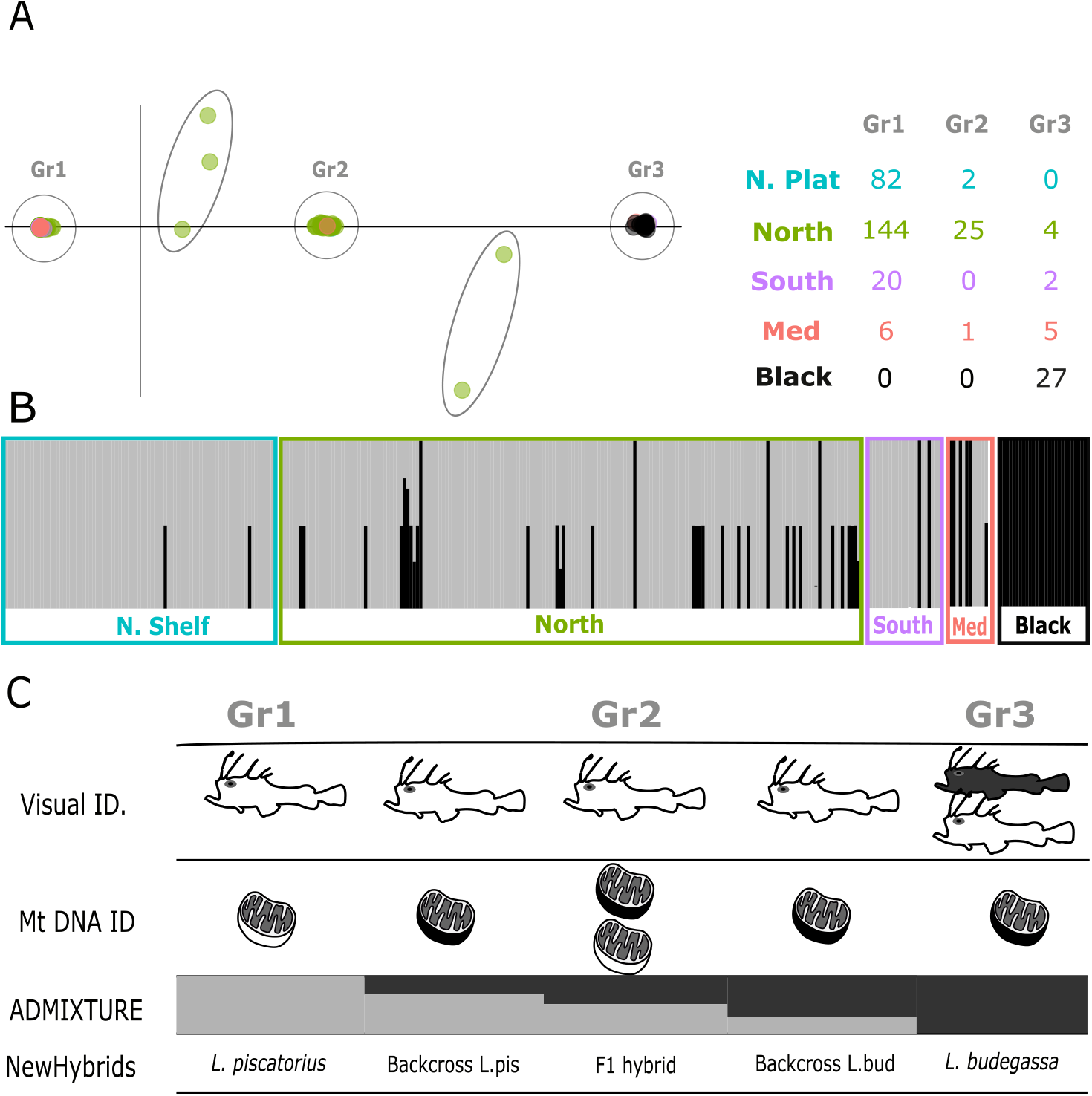
A. Principal Component Analysis (PC1=80.82%; PC2=0.26%) where main differentiated groups are circled. B. Individual ancestry proportions estimated by ADMIXTURE when assuming two ancestral populations. C. For each group identified int the PCA, whether individuals composing it are assigned to L. piscatorius (white fish or mitochondrion), L. budegassa (black fish or mitochondrion) or both according to visual or mitochondrial DNA based species identification, admixture pattern and assignment according to NewHybrids.

### Misidentification and hybridization

Together, these results indicate that some of the samples morphologically identified as *L. piscatorius* are *L. budegassa* despite presenting a white peritoneum (Figure 3C). Samples in the middle group and adjacent ones have a significantly higher observed average heterozygosity (0.77) than samples in first and third groups (0.05 and 0.03, respectively), which could indicate that these individuals are hybrids. This was con-firmed by NewHybrids, which assigned all individuals in the middle group as new gen-eration hybrids between the two species (F1), and the five remaining individuals as backcrosses between F1 hybrids and white or black anglerfish (Figure 3C). Mitochon-drial DNA from all hybrids except for 3 F1 was of *L. budegassa*. Considering the propor-tion of hybrid individuals within the ones bearing *L. piscatorius* or *L. budegassa* mito-chondrial DNA as estimated from RAD-sequencing data, we calculated the percentage of hybrid and misidentified individuals among those provided as *L. piscatorius*. Overall, we found about 4 and 10% misidentified and hybrid individuals, respectively. However, these were not distributed homogeneously across stocks and areas within the same stock (Figure 4). Misidentified individuals were more present in the most southern re-gion of the Southern stock (Portuguese coast) and Mediterranean Sea, with 1/3 of the individuals identified as *L. piscatorius* being *L. budegassa*, less frequent in the Northern stock (2%), and absent in Northern Shelf stock. Hybrid individuals were absent in the Southern stock and North Sea (belonging to the Northern Shelf stock) and were most frequent in the northern Bay of Biscay and Celtic seas (13%).

**Figure 4.**
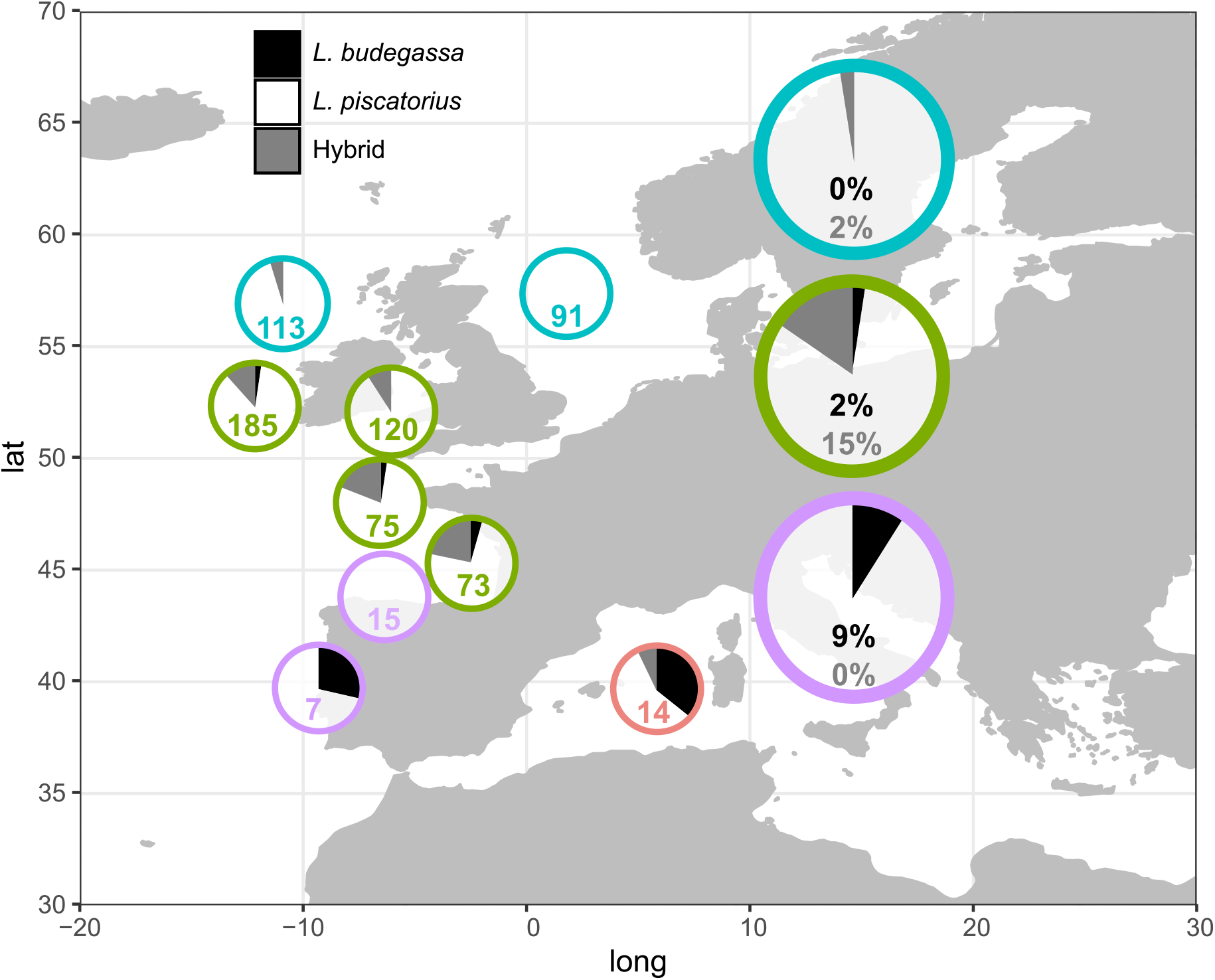
Proportion of hybrid (grey areas and numbers) and *L. budegassa* (black areas and numbers) individuals among those morphologically identified as *L. piscatorius* for each stock (large circles) and, within stocks, for smaller areas (small circles indicating number of sampled individuals per area). Colours refer to stocks as in Figure 1.

### Connectivity of white anglerfish within the Atlantic

The genotype table including only those individuals genetically identified as *L. piscatorius* include 238 or 232 individuals and 6233 or 6246 SNPs when including or not Mediterranean samples, respectively. PCA and ADMIXTURE analyses based on those datasets (Figure 5) reveal strong differentiation between Mediterranean and Atlantic samples (F_ST_ = 0.057) and no genetic differentiation within the Atlantic (F_ST_ = 0), as also suggested by ADMIXTURE (best K=1).

**Figure 5.**
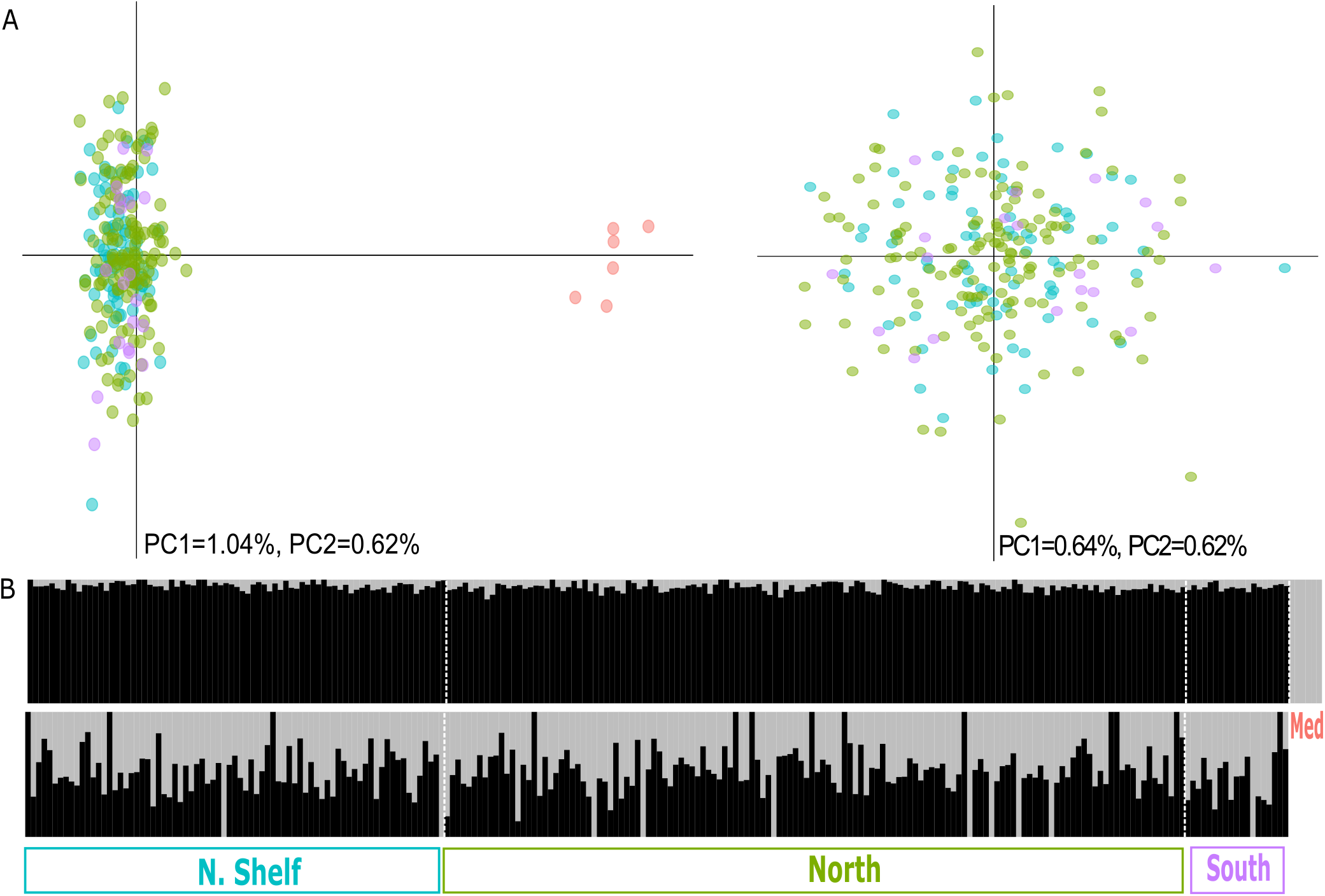
A. Principal Component Analysis (PCA) of samples genetically identified as white anglerfish when including (left) or not (right) Mediterranean samples. B. Individual ancestry proportions of samples genetically identified as white anglerfish estimated by ADMIXTURE when assuming two ancestral populations including (above) or not (below) Mediterranean samples.

## Discussion

### What’s in a white anglerfish sample?

We found that several of the samples provided as white anglerfish (*Lophius piscatorius*) by expert scientists involved in surveys targeting anglerfish were indeed black anglerfish or hybrids. This implies that the commonly used diagnostic character for species identification, the color of the peritoneum, is not discriminative as all the black anglerfish and hybrid individuals provided as white anglerfish had white peritoneum. Indeed, it has already been suggested that some young individuals cannot be distinguished by the color of the peritoneum (*e*.*g*. some that show a white peritoneum with small black dots can be assigned to either black or white anglerfish when using another diagnostic character such as the number of dorsal fin rays (J. Landa and A. Antolínez, unpublished data). Here, we found misidentified individuals that are large (up to 75cm), suggesting that the reason for misidentification is not related to the size of the specimens. In light of these findings, alternative characteristics for species identification are needed (e.g. dorsal and anal fin ray counts or length of the cephalic dorsal fin spines (Caruso, 1986)), so that misidentification does not affect data collection. Additionally, a way of identifying hybrids would be needed. Notably, in our dataset, the hybrid size distributions was significantly smaller than that of the *L. piscatorius* individuals (p<0.05 in the Mann-Whitney test), although the small sample size of hybrids and confounding factors (e.g. different proportion of hybrids in the different stocks, age classes) prevent drawing conclusions from this fact at this stage.

### Causes and consequences of interspecific hybridization

We provide, to the best of our knowledge, the first evidence of natural hybridization between *Lophius piscatorius* and *L. budegassa*, a phenomenon that can occur between closely related species sharing morphological, ecological and reproductive compatibilities (Montanari, Hobbs, Pratchett, Bay, & Van Herwerden, 2014). *Lophius spp*. hybrids were predominant in Northern Bay of Biscay and Celtic seas, where both species’ distribution range overlap and have been historically abundant (Figure S2).

There is also overlap in both species’ bathymetric distribution (20-1000m and 100-500m depth respectively for white and black anglerfish)(Azevedo, 1995; Caruso, 1986; García-Rodríguez, Pereda, Landa, & Esteban, 2005; Quincoces, Santurtún, & Lucio, 1998), and in spawning periods, both spawning in winter and spring (Ofstad, Angus, Pedersen, & Steingrund, 2013; Quincoces, 2002.). This overlap in space, depth and time could have even been increased by recent changes of geographical and bathymetric distributions of both species, potentially as a consequence of climate change (Maltby, Rutterford, Tinker, Genner, & Simpson, 2020). According to mitochondrial DNA, which is maternally inherited in most animals (Moritz, Dowling, & Brown, 1987), we found that most hybrids (27 out of 30) resulted from a black anglerfish mother and a white anglerfish father. We found a few backcrosses (hybrids crossed with parental species) and no crosses involving two hybrids. This could be due the presence of a stable hybrid zone with hybrids being less able to produce offspring (Hayden et al., 2010; Mirimin et al., 2014) or to recent changes that have induced hybridization between both species so newly that they did not have time to produce a hybrid-derived population. The presence of a stable hybrid zone between black and white anglerfish does not forcedly imply changes in the evolution of the parental species but could imply management uncertainties. However, if hybridization is recent, we cannot discard a process of evolutionary novelty (Budd & Pandolfi, 2010), through parental species acquiring new functions (T. M. Anderson et al., 2009) and even producing new species (Verheyen, Salzburger, Snoeks, & Meyer, 2003), or of biodiversity loss, through extinction of parental species (Seehausen, 2006). Thus, monitoring the hybrid zone and adjacent areas is crucial to understand the role of hybridization in management and conservation of white and black anglerfish.

### Management implications of stock connectivity, hybridization and misidentification

Our results show that the Northeast Atlantic and Mediterranean white anglerfish populations are genetically isolated, challenging previous findings based on genetic data (Charrier et al., 2006), and that the white anglerfish within the Northeast Atlantic Ocean constitutes a genetically homogeneous population, shedding light into previous unconclusive results based on genetic and non-genetic data (Blanco et al., 2008; Cañás et al., 2012; Crozier, 1987; Charrier et al., 2006; J. Landa et al., 2008; Laurenson et al., 2005). These results point towards the necessity to harmonize current stock definitions within the Northeast Atlantic. Morphometric characteristics such as length of maturity (Duarte, Azevedo, Landa, & Pereda, 2001) or weight-length relationships (Jorge Landa & Antolínez, 2018) differ between stocks. These differences are potentially due to sampling biases, differences in scales and/or measurement methods or interannual differences among studies, as well as the possible influence of different fishing pressure or environmental conditions between areas. However, they could also be due to the inclusion of different proportions of misidentified and hybrid individuals in each stock.

Although in the Northern and Southern stocks the assessment is done separately for white and black anglerfish, they are combined for management, being Total Allowable Catches (TAC) for *Lophius spp*. In the Northern Shelf stock, the two species are assessed and managed as one assuming that black anglerfish in this area are rare.

However, the obvious increase of black anglerfish in this stock (Figure S2) has created concern and suggestions to separate species have been made (ICES, 2018). The deliberate (in the North Shelf stock) or non-deliberate (in the Northern and Southern stocks) inclusion of an unknown (and perhaps variable) proportion of black anglerfish in the white anglerfish assessment will likely lead to some bias. Therefore, some corrections could be applied to the total catches and the length composition data.

However, the proportions of black anglerfish included in these assessments seem relatively small (<10%) and would likely not have a major impact on the assessments. The implications of including hybrids in the stock assessment are more difficult to anticipate as biological characteristics of hybrids are unknown. The main concern is that inclusion of hybrids would lead to erroneous inferences of reproductive potential because this feature is inferred from the biomass of mature fish and the reproductive output of anglerfish hybrids is unknown. If, as in many other species (Mallet, 2007), they have no or neglectable offspring, they should not be included in the spawning biomass as otherwise they could lead to an overestimation of productivity, even to a level below of required to support a sustainable fishery (Morgan et al., 2012). On the other hand, if, as the presence of backcrossed individuals suggest, they are reproductively viable, further studies are required to assess hybrid fitness compared to pure individuals.

Despite supporting stock merging, the analyses presented here suggest a series of concerns that should be considered for white anglerfish management and which affect differently each of the stocks. The southernmost locations of the Southern Stock and the Mediterranean Sea are more affected by misidentification, the northernmost locations of the Northern stock are more affected by hybridization, and the North Shelf stock does not seem to be affected by either. The over and under representation of misidentified individuals (black anglerfish with white peritoneum) in the south and north respectively could be simply explained by differences in black anglerfish abundance. However, both species are moving northwards, as shown in the species distribution maps (FigureS2), and so the proportion of misidentified individuals could increase in northern areas as black anglerfish becomes more abundant. This northward movement of both species could also contribute to enlarge the hybrid zone towards northern areas. If the proportion of hybridization and mislabeling varies significantly over time, then this will have implications for the robustness of the assessment for these stocks. Thus, genetic studies monitoring white and black anglerfish populations across the Northeast Atlantic and Mediterranean Sea through time are needed.

### Outlook

Our study shows that genetic analyses can be used to confirm or reject existing hypothesis about, for example, stock connectivity but, most importantly, that they can reveal hidden phenomena that were not foreseen, such as the thus far unknown hybridization between white and black anglerfish. Yet, despite the power of genetics to provide fisheries assessment relevant information, there are still barriers for the uptake of genetic data by fisheries management, which can be due to a variety of factors (Bernatchez et al., 2017; Ovenden, Berry, Welch, Buckworth, & Dichmont, 2015), such as lack of clear communication of genetic concepts by geneticists to end-users and reluctance to change or adapt established assessment and management procedures. The white anglerfish is a highly valuable species (around 30K tons with a corresponding value of around 142 million euro; data extracted from https://stecf.jrc.ec.europa.eu/dd/fdi) with a well-established data collection, and assessment and management frameworks. Thus, we anticipate that these results will set the basics for an imminent genetics informed fisheries management for this species. Indeed, a genetics informed fisheries management is essential to ensure the basics of fisheries science, whereby maximum sustainable yield can only be reached by efficient management of distinct populations.

## Supporting information

Suppementary Material

## Acknowledgements

We would like to thank Arkaitz Pedrajas and Arnaitz Mugerza (AZTI), Liz Clarke, Jim Drewery and Ruadhan Gillespie-Mules (MSS, Scotland), Eoghan Kelly and David Tully (MI, Ireland), Jose Carlos Fernández Franco, Ángela Cortina Burgueño and Oscar Fernández Acevedo (OPPF-4, Spain), the fisheries surveys team (CEFAS), Izaskun Preciado and Susana Ruiz (IEO), Corina Chaves (IPMA, Portugal), Anabel Colmenero (ICM-CSIC, Spain), Chrysoula Gubili (FRI, Greece) and Nota Peristeraki (HCRM, Greece) for sampling and Elisabete Bilbao for laboratory work. This work has been funded by the Joint Research Centre (European Commission), through the project GECKA (Contract JRC/IPR/2017/D.2/0016/NC), the Department of Environment, Planning, Agriculture and Fisheries (Basque Government), through the project GENGES and a predoctoral grant to Imanol Aguirre-Sarabia, and by the Department of Education (Basque Government), through a predoctoral grant to Iker Pereda. This is contribution 1043 from the Marine Research Division of AZTI.

## Data availability statement

Demultiplexed and quality-filtered RAD-tag reads are available at the NCBI Sequence Read Archive (SRA) under accession number SUB9963797 and final datasets and scripts utilized for this project are available at https://github.com/rodriguez-ezpeleta/Lophius_PopGen

## Author Contributions

NRE, JTM, AZ, MS, IQ and IC designed the study. HG, FB, IH and JL provided samples. IM generated data. IAS, NDA, IPA and NRE analyzed data. IAS, NRE, AU, HDG, FB, IH, JL, IQ interpreted data. IAS and NRE wrote the first draft of the manuscript and integrated input from all authors.

