## Supplementary material for "Evidence of stock connectivity, hybridization and misidentification in white anglerfish support the need of a genetics-informed fisheries management framework": Suppementary Material

### Supplementary Tables

**Table S1.** Samples used for this study with collection data (survey, responsible institute, date, location), biological variables measured (morphological identification, depth in meters, length in cm, weight in g, sex and maturity) and molecular analysis information (used for RAD-seq or not, replicate or not, genetic identification and hybrid class).

[attached excel file]

**Table S2.** Number of individuals and SNPs including in the catalog and remaining after each filtering steps (genotyping rate higher than 0.95 and minimum allele frequency (MAF) bigger than 0.05) for each dataset (All: all individuals included; L. pis: only those individuals genetically identified as *L. piscatorius*; L. pis without MED: L. pis dataset without Mediterranean Sea individuals).

| | **All** | | | | **L. pis** | | **L. pis without MED** | |
|---|---|---|---|---|---|---|---|---|
| | **de novo** | | **Ref. mapped** | | **de novo** | | **de novo** | |
| | **Inds** | **Tags/SNPs** | **Inds** | **Tags/SNPs** | **Inds** | **Tags/SNPs** | **Inds** | **Tags/SNPs** |
| **Catalog** | 327 | 25892 | 326 | 35561 | 241 | 23301 | 235 | 23095 |
| **Genotyping rate** | 323 | 24346 | 326 | 31545 | 238 | 21705 | 232 | 21579 |
| **MAF** | 323 | 16712 | 326 | 23126 | 238 | 6233 | 232 | 6246 |

**Supplementary Figures**

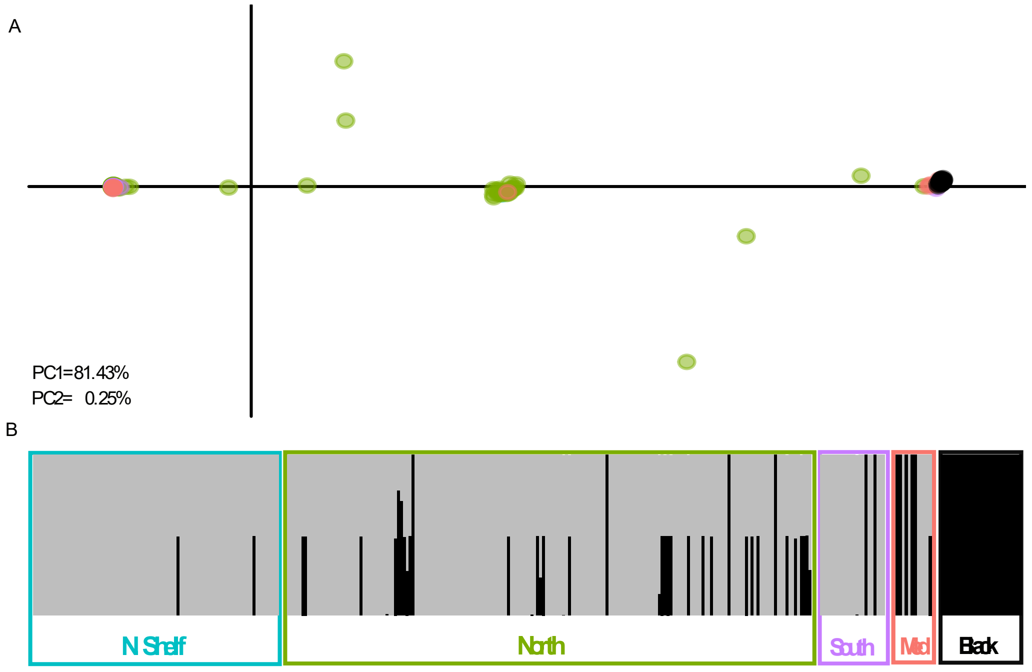


Figure S1. A. Principal Component Analysis (PCA) for *L. piscatorius* and for *L. budegassa* samples coloured according to sample location (see panel B). B. Individual ancestry proportions estimated by ADMIXTURE when assuming two ancestral populations. These results are derived from the reference mapped catalog.

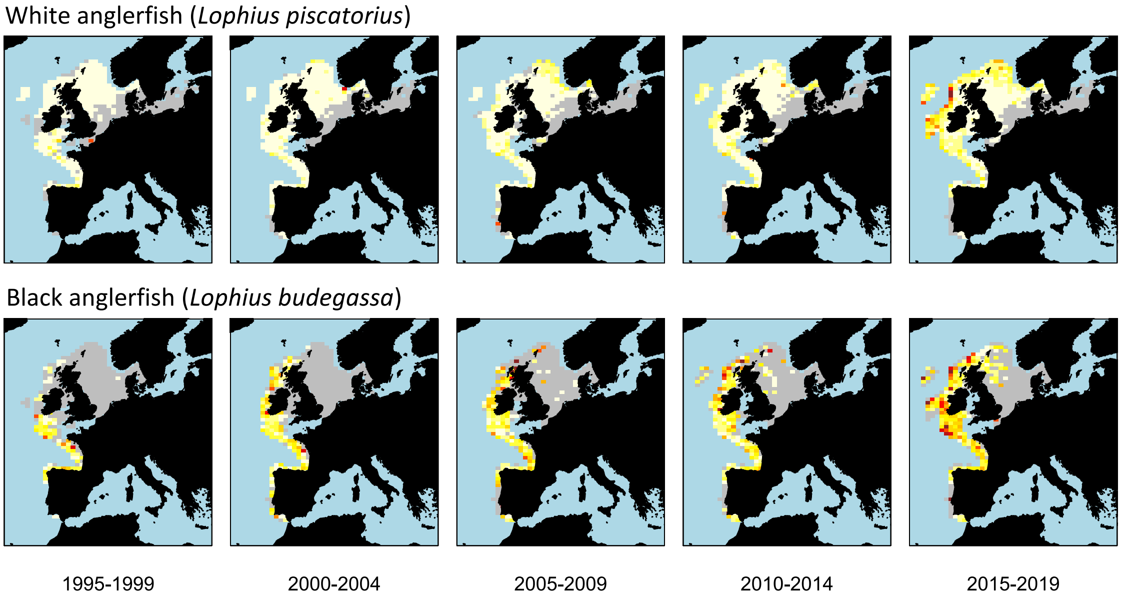


Figure S2. Maps showing catch per trawl time for white (top) and black (bottom) anglerfish considering trawl surveys targeting anglerfish (BITS, NS-IBTS, SWC-IBTS, EVHOE, SP-NORTH, SP-ARSA, ROCKALL, PT-IBTS, IE-IGFS, NIGFS, DWS, SCOROC, SCOWCGFS, IE-IAMS, FR-CGFS). Using information for each haul (latitude, longitude, haul duration and catch weight per species), obtained from the icesDatras R library (https://github.com/ices-tools-prod/icesDatras) data was aggregated by five consecutive years and ICES statistical rectangles and converted to grams per minute (g/min) per species. For those hauls with catch data as number of individuals instead of weight, weight was estimated using weight-length relationships. For each five-year period and rectangle, g/min per species was plotted in a gradient from white (minimum abundance) to red (maximum abundance, which was 2053 and 335 for white and black anglerfish, respectively) using grey to represent areas where sampling was performed but no catch was reported.
